# Mothers respond to biological pup calls with heart rate changes in Japanese house bats, *Pipistrellus abramus*

**DOI:** 10.1101/2025.07.21.666052

**Authors:** Kazuki Yoshino-Hashizawa, Midori Hiragochi, Kohta I Kobayasi, Shizuko Hiryu

## Abstract

Maternal care is essential for offspring survival in mammals, especially in colonial species where mothers must recognize their own young among many. In the Japanese house bat, *Pipistrellus abramus*, mothers identify their pups using acoustic cues, particularly isolation calls (ICs) produced by newborns. However, the physiological mechanisms underlying such maternal recognition remain largely unknown. Here, we investigated maternal emotional responses of mother bats to pup calls by measuring heart rate (HR) changes during controlled playback experiments. We recorded ICs from 2-day-old pups and echolocation calls (ECs) from 30-day-old pups, then presented these sounds to mothers after their pups had become independent. HR significantly increased in response to calls from the mothers’ own pups—both ICs and ECs—but not to calls from non-family pups. Among all stimuli, ECs from their own older pups evoked the largest HR increase, indicating strong physiological arousal and suggesting sustained maternal responsiveness despite developmental changes in call structure. In contrast, ECs from an unfamiliar adult also induced HR elevation, possibly reflecting general social arousal rather than maternal recognition. These findings demonstrate that *P. abramus* mothers exhibit selective physiological arousal to their own pups’ vocalizations, and that HR provides a sensitive physiological index of maternal motivation and recognition based on dynamic acoustic information.

## Introduction

Maternal lactation is a fundamental behavior in mammals, providing offspring with essential nutrients required for survival and proper development during the early stages of life. If a female were to nurse the offspring of another species or unrelated individuals, her ability to provide sufficient milk to her own offspring would be compromised, ultimately reducing her reproductive fitness. Consequently, through long-term adaptive evolution, adult female mammals have developed the ability to accurately recognize their own offspring [1]. Mammalian females utilize multiple sensory modalities for offspring recognition, including visual [2], olfactory [3], and auditory [4] cues, or a combination of these [5]. Among these, acoustic cues are particularly important in low-visibility environments and are widely used by many animals, especially nocturnal species. Acoustic signals emitted from pups, including isolation calls (ICs), are recognized by their mothers [6–9]. ICs are often perceived as distress calls, reflecting the negative emotions states of offspring [10], and play critical role in mother-pup communication and individual recognition, particularly when pups are separated from their mothers [11]. Most research on mother-offspring communication via ICs has focused on ultrasonic vocalizations (USVs) in rodents, commonly used as experimental models due to their versatility [12]. Maternal behavior tests in rodents have demonstrated that mothers exhibit nurturing behaviors – such as pup retrieval-, immediately after locating their pups via USVs. However, mice and certain other rodents are known to care not only for their own pups but also for unrelated pups, primarily relying on olfactory cues for pup recognition [13–15].

In contrast, echolocating bats present a unique model for studying mother-offspring communication based on acoustic cures. It has been reported that bat pups produce ICs, and mother bats can discriminate own pups using these calls [16–21]. ICs in bats are typically characterized by long duration, low frequency, and a high repetition rate, allowing mothers to identify their own pups even within large colonies [17,18,20]. Moreover, despite developmental changes in call structure, mothers may maintain the ability to recognize their pups throughout the nursery period [22]. In addition to ICs, echolocation calls (ECs) may also contribute to individual identification. It has been theorized that ECs in some bat species may encode information of sex [23,24], social affiliation [25], and individual identity [24,26], with receivers capable of decoding these information. Collectively, these findings suggest that mother bats are well-suited model animals for studying maternal responses to pup vocalizations, particularly in the context of individual recognition.

While behavioral responses, such as pup retrieval, have been widely used to infer maternal recognition, such behaviors do not fully capture the underlying emotional and motivational states of the mothers. Previous studies have suggested that pup USVs elicit nurturing behaviors via emotional arousal in rodents [12,27,28]; however, arousal is not always apparent through behavior alone. To assess internal emotional states directly, physiological measures offer a valuable complementary approach. Heart rate (HR) is widely used as a minimally invasive physiological indicator of emotional arousal across various species. Monitoring HR can reveal subtle internal state changes that may not be observable through behavior, providing insight into the affective component of maternal responses. HR measurements in bats may help clarify whether mothers exhibit specific physiological responses to their own pups’ calls, or whether all ICs are perceived as signals of negative emotional valence. In this study, we measured HR responses in mother bats during playback of pup vocalizations to investigate their emotional reactions. Building on these approaches, our study aims to explore maternal emotions and motivation in bats from a psychophysiological perspective by examining HR responses to acoustic stimuli, with a particular focus on the discrimination of calls from their own versus non-family pups.

## Materials and Methods

### Study species and captive status

The target species was the Japanese house bat *Pipistrellus abramus* (Temminck, 1840), an insectivorous, aerial-hunting species belonging to the bat family Vespertilionidae. For echolocation, adults of this species use frequency-modulated (FM) pulses that sweep downward from initial frequencies around 80 kHz to terminal frequencies (TF) of around 40 kHz [29,30]. Their auditory sensitivity exhibits a broad U-shaped curve in the 4-80 kHz frequency range, with low thresholds between 20 and 50 kHz and the highest sensitivity in the 35-50 kHz range [31]. Vocalizations in *P. abramus* pups differ from adult echolocation calls and appear to serve as precursors of echolocation calls as well as ICs [22]. They raises their pups in colonies of 50–200 individuals, where multiple families of females and their pups coexist [32].

In May 2022, four late-pregnancy female bats were captured from a colony roosting in bridge girders near the campus of Doshisha University, in Kyotanabe, Kyoto, Japan. Each female was individually banded with a marked ring on her forearms (2.0 mm internal diameter, 4.0 mm height, Porzana Ltd, Winchelsea, U.K.). All four females gave birth to litters of three pups in the laboratory over three days, from 20 to 22 June 2022. We checked the condition of the pups daily, and they were well cared for by their mothers. The pups’ body mass ranged from 1.1 to 1.3 g, whereas adult body mass ranged from 5 to 8 g. Since two pups died shortly after birth (pup-Ba from mother-B and pup-Db from mother-D), we recorded the vocalizations of the ten surviving captive-born pups (three female and seven male) from the four mothers during their first postnatal month. The pups were identified as follows: pup-Aa, -Ab, and -Ac from mother-A; pup-Bb and -Bc from mother-B, pup-Ca, -Cb, and -Cc from mother-C; pup-Da and -Dc from mother-D. Each pup was marked using a harmless animal marker (FG2200G, EXEC, Japan) applied to a small area of skin on its back. All mothers and pups were kept in a rearing cage (25 cm × 17 cm × 17 cm) under controlled conditions, including a regulated temperature (25 ± 2°C), humidity (55 ± 5%), and light–dark cycles following the natural photoperiod in Kyotanabe, Kyoto, Japan. The female bats were fed mealworms, provided with water, and given an adequate amount of vitamins to ensure proper nutrition.

### Experiment I: Recording vocalization from pups

Pup vocalizations were recorded daily during their first post-natal month. To record each pup vocalizations, one pup was gently separated from its mother and put on an echo-soundproof sponge (30 cm× 30 cm) in the experimental room. After the recording, the pups were returned to their mother, and the same procedure was repeated for its siblings. Vocalizations were recorded using an ultrasonic microphone (Ultrasoundgate CM16/CMPA, Avisoft Bioacoustics, Germany) placed at approximately 20 cm from the pup for 3 minutes with a sampling frequency of 250 kHz and a resolution of 16 bits. As a control, we also recorded vocalizations from one adult male bat using same recording setup. A custom MATLAB program was used to analyze the acoustic characteristics of the calls, including duration, terminal frequency (TF), and initial frequency (IF), obtained from the spectrogram at −20 dB relative to the peak intensity of the call. These characteristics were then used to classify the calls into the ICs and echolocation calls (ECs), based on previous study [22].

### Experiment II: ECG recording from mother bats

To avoid interference with maternal care, electrocardiograms of four mother bats were recorded from 38 to 45 days after delivery, when the pups had matured. During these recordings, sound stimuli were presented. All sound stimuli were taken from pup vocalizations recorded in Experiment I, specifically from males at 2 days after birth (DAB) for ICs and at 30 DAB for ECs, and were normalized to a length of 3 seconds, containing between 15 to 33 calls per stimulus.

Five types of sound stimuli were presented to each mother bat: (ICo) ICs from their own pups at 2 DAB, (ECo) ECs from their own pups at 30 DAB, (ICn) ICs from non-family pups at 2 DAB, (ECn) ECs from non-family pups at 30 DAB, and (ECu) ECs from an unfamiliar adult bat. For stimuli ICn and ECn, two or three non-family pup calls were uses, consisting of calls from pups of different mothers (e.g., mother A was exposed pup Bc’s and Cc’s calls for both stimuli ICn and ECu). All stimuli were normalized the stimuli based on peak-to-peak amplitude, as it was difficult to control the loudness due to differences in call duration and frequency range of ECs and ICs.

This experiment was carried out in a soundproof chamber (1.4 m×1.4 m×2.1 m). Each subject was positioned in the center of a circular table using soft sponges for stabilization. Sound stimuli were presented at 100 dB peak-to-peak SPL via a tweeter (PT-R7III, Pioneer, Japan) located approximately 15 cm from the subject. The stimuli were played through an audio interface (Rubix44, Roland, Japan) at a sampling frequency of 192 kHz and a resolution of 16 bits, amplified by an amplifier (STR-DH190, Sony, Japan) connected to the tweeter.

A total 39 sound stimuli were presented to each mother over two days, with each of the five stimulus types presented at least six times. One of the four mothers died before the second day of the experiment and was therefore exposed to only 18 stimuli. The order of stimulus presentation was randomized, except for the first stimulus of each day. To avoid potential startle responses from the first stimulus, the adult EC was always presented first and was excluded from subsequent analysis.

ECG electrodes were attached subcutaneously in a Lead II configuration at three positions: left leg (positive), right thumb (negative), and right leg (reference). Lead II is known to provide a highest signal-to-noise ratio for ECG recording in echolocating bats [33]. Subjects were anesthetized with 2–4% isoflurane (Pfizer, USA) during surgery. Fur was shaved at three dorsal sites (base of the right arm, left leg, and right leg), and 2% xylocaine jelly (Sandoz, Switzerland) was applied as a local anesthetic to facilitate the subcutaneous insertion of small safety-pin electrodes. Electrodes were connected to the ECG recording system (C3410 / C3314 / C3216 / C3100, Intan technology, USA) and, signals were recorded using vendor software (RHX software, Intan technology, USA) at a sampling rate of 30 kHz. ECG recordings were obtained by measuring the potential differences between the positive and the negative electrodes. Before presenting sound stimuli, subjects were allowed to acclimate until their HR stabilized at approximately 200-300 beats per minute (bpm). ECGs were recorded from 50 seconds before to 100 seconds after the onset of each sound stimulus (one trial). After each presentation, we ensured that the HR returned to baseline before presenting the next stimulus.

### ECG processing and statistical analysis

All digital signal processing was performed using a custom MATLAB program (R2021a, MathWorks, USA). To reduce noise, a 1–650 Hz band-pass filter was applied to the raw ECG. Instantaneous HR was calculated at the reciprocal of the R-R interval, which was automatically detected from the filtered ECG signal. HR values were then binned into one-second intervals, and relative HR changes were calculated by subtracting the average HR in −10 to 0 second period as the baseline. HR increases were quantified by averaging HR values over the 0–30 second period following the onset of each sound stimulus type [34].

All statistical analyzes were performed using a custom MATLAB program (R2024a, MathWorks, USA). To test for significant effect of factors (relationships: biological or non-biological family, and vocalization type: IC or EC), we applied a two-way ANOVA test to average HR values over the 0–30 second period after the stimulus onset in Experiment II excluding adult EC (stimulus ECu). Finally, to discuss HR increases to each acoustic stimulus, we applied a one-sample Wilcoxon signed-rank test with Bonferroni correction to average HR values over the 0–30 second period after the stimulus onset in Experiment II including adult EC (stimulus ECu).

## Results

### Experiment I: Developmental changes in ultrasonic vocalizations by pups

All pups vocalized ICs until approximately 10 to 12 DAB (Fig. 1 top). Although pups already vocalized ICs at 0 DAB, their amplitude was unstable, making it difficult to evaluate their acoustic components. At 2 DAB, the fundamental frequency component of ICs consisted of FM sounds that descended from the initial frequency (IF) of the first harmonic, approximately 35 kHz, to the terminal frequency (TF), around 20 kHz, across all pups (see Table S1). As the pups developed, both IF and TF increased, accompanied by a reduction in call duration. By 10 DAB, ICs reached approximately 40 kHz (IF), 25 kHz (TF) and a duration of 10 ms.

**Fig. 1.**
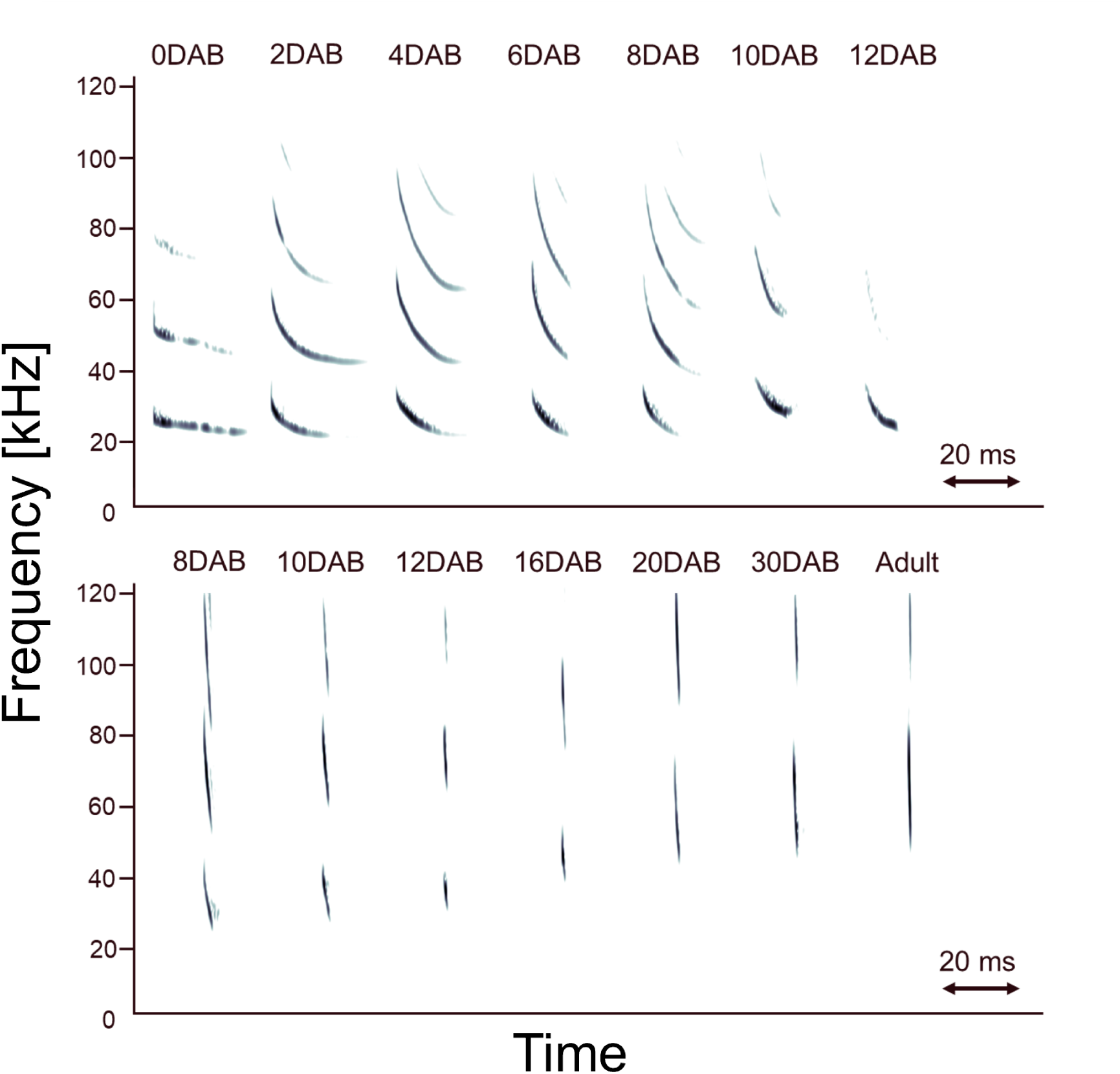
Developmental changes in ultrasonic vocalizations by pups of *P. abramus*. Spectrograms of ultrasound vocalizations produced by a pup. Upper row: isolation calls (ICs) recorded when the pup was separated from its mother, from 0 to 12 DAB. Bottom row: echolocation calls (ECs) recorded when the pup was actively exploring from 8 to 30 DAB, along with a representative EC from an adult bat.

In contrast, ECs first appeared at approximately 8 DAB (Fig. 1 bottom). At this stage, the IF of the first harmonic was around 50 kHz, and the TF was around 30 kHz. Both frequencies increased with postnatal development, while call duration gradually shortened, reaching levels comparable to those of adult ECs by 30 DAB (Table S1). Figure 2A shows an example of postnatal vocalization development in a representative pup (Ab). Throughout this developmental process, the pups’ vocalizations progressively approached the acoustic characteristics of adult ECs, a pattern consistently observed in other pups as well (Fig. 2B).

**Fig. 2.**
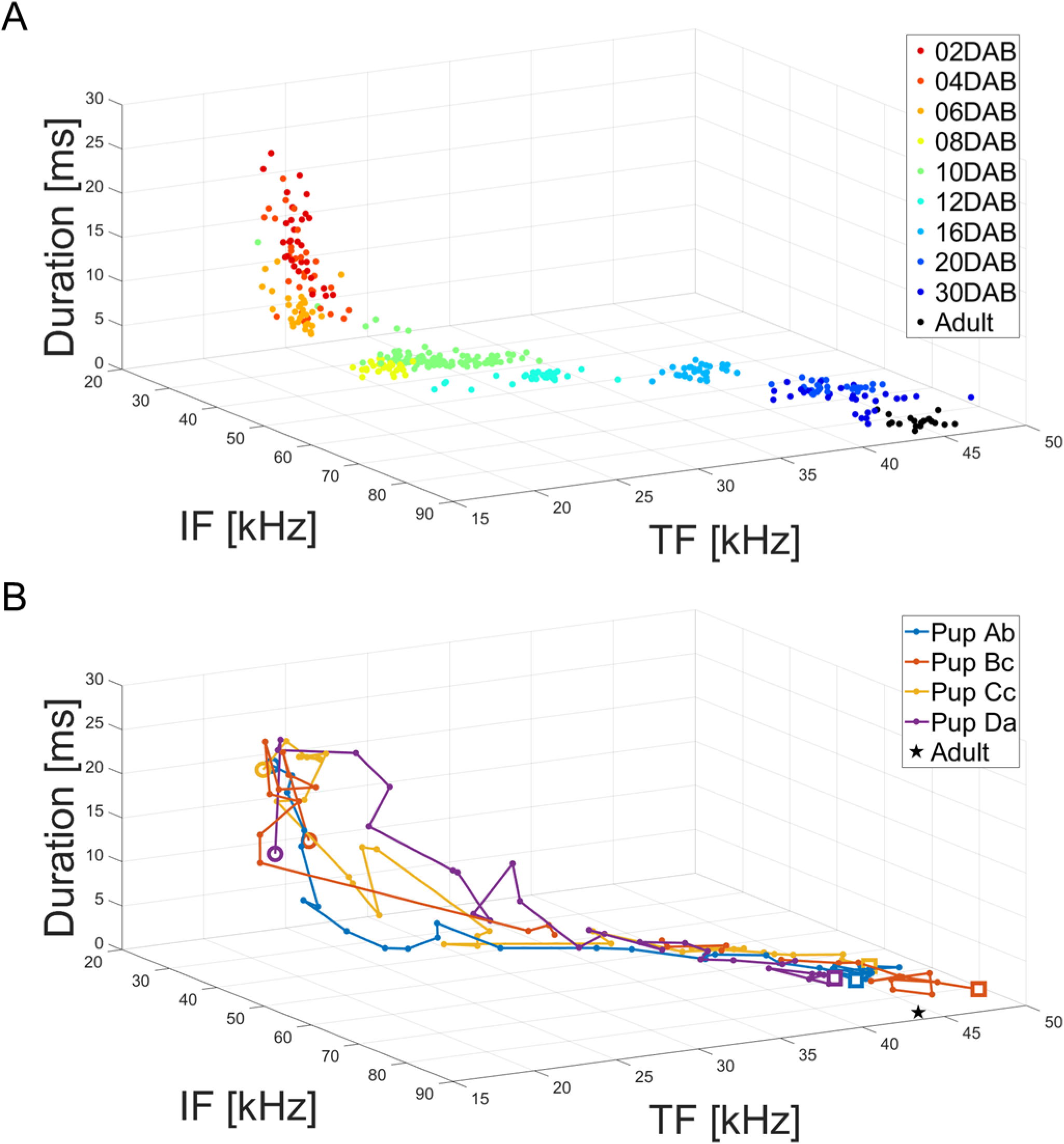
Developmental changes in ultrasonic vocalizations by pups of *P. abramus*. (A) Example of developmental changes in duration, initial frequency (IF) and terminal frequency (TF) from 0 to 30 DAB in a representative pup (Ab). Red to blue points indicate changes over time, and black points represent ECs from an adult bat. (B) Average developmental changes in vocalization parameters across pups from different mothers, along with the adult bat EC (black star marker). The large open circle marker indicates vocalizations at 0 DAB, and the large open square marker indicates those at 30 DAB.

### Experiment II: Mother bat’s ECG changes for sound stimuli

As illustrated in Fig. 3, the ECG and HR responses of the mother bats were recorded before and after the presentation of sound stimuli. The ECG signals were recorded with a sufficient signal-to-noise ratio to allow clear identification of R waves for HR calculation (Fig. 3A). Across all five stimulus types, the mean HR showed an immediate increase following stimulus onset (Fig. 3B).

**Fig. 3.**
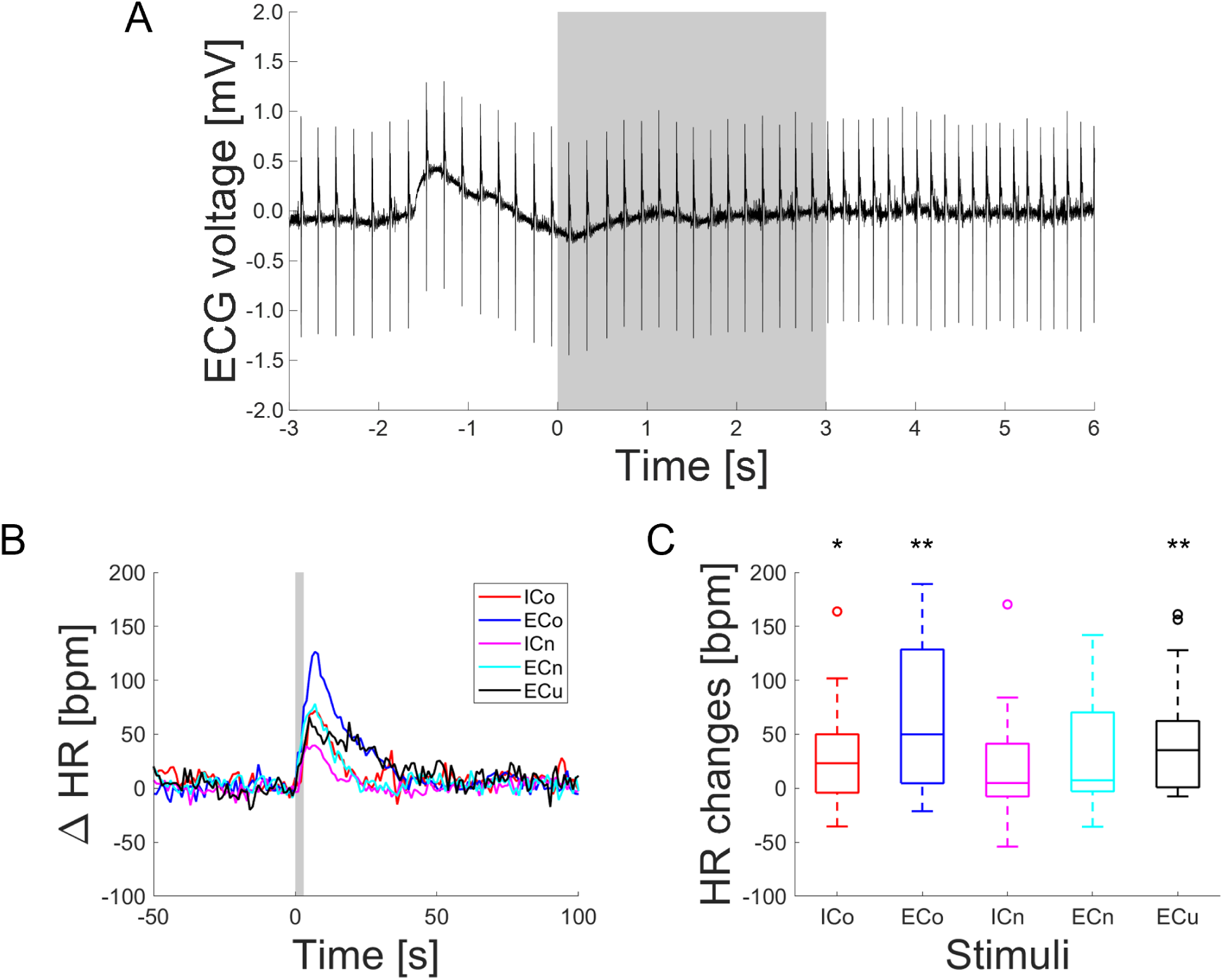
Heart rate responses to acoustic presentation in mother bats. (A) Example of a raw electrocardiogram (ECG) recording from a mother bat. The gray area indicates the period of acoustic stimulus presentation with echolocation calls (ECs) from the mother’s own pup (Stimulus ECo). (B) Average heart rate (HR) responses of all mothers to acoustic stimuli. The baseline was calculated as the average HR during the period from −10 to 0 seconds before stimulus onset. The five stimulus types were: (ICo) isolation calls (ICs) from their own pups at 2 DAB, (ECo) ECs from their own pups at 30 DAB, (ICn) ICs from non-family pups at 2 DAB, (ECn) ECs from non-family pups at 30 DAB, and (ECu) ECs from an unfamiliar adult bat. (C) Distribution of HR increases from 0 to 30 seconds after stimulus onset. Each box plot shows the median (red line), interquartile range (box edges), and the minimum and maximum values (whiskers). Statistical significance is indicated by \**P*<0.05, \*\**P*<0.01 (Wilcoxon sign rank test with Bonferroni correction).

A two-way ANOVA reveals significant main effects of relationships—that is, whether the presented calls were from the mother’s own pups (kin) or from non-family pups (non-kin)—(*F*_(1,110)_ = 4.72, *P* = 0.032), and vocalization types (ICs or ECs) (*F*_(1,110)_ = 7.16, *P* = 0.009), but no significant interaction between these factors (*F*_(1,110)_ = 0.61, *P* = 0.435). The largest increase in means HR was observed in response to ECs from their own pups at 30 DAB (ECo), reaching approximately 130 bpm at its peak, which was significantly higher than baseline (*P* = 0.001; Fig. 3C). ECs from an unfamiliar adult bat (ECu) also elicited a significant increase in HR (*P* = 0.001). In response to ICs from their own pups at 2 DAB (ICo), HR increased by approximately 70 bpm (Fig. 3B), which was likewise significantly higher than baseline (*P* =0.040). As shown in Fig. 3C, although ICs from non-family pups at 2 DAB (ICn) and ECs from non-family pups at 30 DAB (ECn) induce slight increases in mean HR, these changes were not statistically significant (ICn: *P* = 0.137, ECn: *P* = 0.171).

## Discussion

In Experiment II, the HR of mother bats significantly changed with the factor of relationship type, categorized as either kin or non-kin (two-way ANOVA test, *P* = 0.032). In addition, the HR of mother bats significantly increased compared with baseline in response to their own pups’ vocalization at both 2 DAB and 30 DAB (stimuli ICo and ECo), whereas calls from non-family pups (stimuli ICn and ECn) did not induce significant changes (Fig. 3C). In *P. pipistrellus*, mothers do not exhibit care toward non-family pups [35]. Thus, the significant HR changes in response to biological pup calls in *P. abramus* may reflect a different level of maternal motivation depending on relatedness. Previous studies in rodents have shown that ICs are associated with distress signals, eliciting arousal or stress responses in mothers [10]. In addition, such recognition of pups elicits HR acceleration [36]. Furthermore, not only social calls that include ICs but also ECs have been shown to encode various types of social information, including sex, individual identity, group membership, and behavioral states in other species [23,24,26,37,38]. Similarly, *P. abramus* may also extract social information from ECs, which could explain the strong HR response to their own pups’ ECs at 30 DAB (stimulus ECo). It is therefore plausible that mothers of Japanese house bats also experience arousal upon hearing their own pups’ calls, both ICs and ECs. This arousal is reflected in physiological changes, specifically increased HR as a result of sympathetic nervous system dominance, then it may promote maternal behaviors.

In addition, the vocalization type factor significantly affected the HR response in mothers (two-way ANOVA, *P* = 0.009). ICs at 2 DAB are thought to play an important role in pup survival and were hypothesized to elicit a stronger maternal response than ECs at 30 DAB. However, our HR results did not support this assumption; rather, ECs elicited a significantly stronger HR increase than ICs (ECs: 50.95±10.30, ICs: 18.25±5.12; mean±s.e.m.). One possible explanation for this discrepancy is that mother bats—as well as other mammals such as goats and sea lions [7,39]—may continuously update their memory of pup calls on a daily basis. In this study, ECG measurements of mother bats began approximately 40 days after delivery (38 to 45 days), when the pups had already become independent. This timing was chosen to minimize stress on lactating mothers and to prevent potential harm to immature pups, which cannot maintain their body temperature for more than an hour. It is therefore possible that the calls at 30 DAB more closely resembled the most recently experienced pup vocalizations, whereas the calls at 2 DAB may have appeared unfamiliar or unnatural during the experiment. As illustrated in Figure 1, the acoustic characteristics of pup vocalizations change daily. We therefore hypothesize that mothers’ recognition abilities may undergo continuous adaptation. To address memory-related effects linked to vocal changes during development, future studies should incorporate experimental paradigms, such as daily auditory playback experiments, to evaluate how dynamic memory updating influences maternal motivation.

Furthermore, the mothers exhibited significant HR increases in response to ECs from an unfamiliar adult bat (stimulus ECu), even though this stimulus was intended as a social control stimulus. Although we could not directly assess the cause of this response, one possibility is that, around 40 days postpartum, when the experiment was conducted, females were not yet in the mating season. Consequently, hearing adult male calls might have triggered a rejection-related physiological response, potentially mediated by the ventromedial hypothalamus and the sympathetic nervous system [40,41]. Further investigation is warranted to clarify the neural mechanisms underlying reproductive status, social cue processing, and vocal communication in mother bats.

The findings of this study demonstrate that maternal internal state changes in response to pup calls can be assessed through HR monitoring. Specifically, HR increased in response to calls from the mothers’ own pups but not to calls from non-family pups, suggesting that HR reliably reflects maternal recognition and motivation in *P. abramus.* Previous studies have reported HR increases in echolocating bats in response to negative social calls [34,42–44], indicating that HR is a useful physiological indicator of emotional arousal elicited by acoustic stimuli. However, a limitation of using HR alone is its inability to distinguish between positive and negative emotional valences, as elevated HR may occur in both distress and affiliative contexts. In other animal species, including humans, heart rate variability (HRV) is commonly used to infer the balance between sympathetic and parasympathetic nervous system activity and to provide a more nuanced assessment of emotional state [45–47]. To evaluate more complex emotional dimensions, such as the strength of mother-offspring bonds, feelings of security, empathy-like responses, and developmental maturity, it will be important to combine HR measurements with HRV analyses. Importantly, HRV has proven particularly effective in detecting subtle emotional responses to social acoustic stimuli, such as vocal communication. Nevertheless, in bats, standard frequency band definitions for low-frequency and high-frequency components, essential for HRV power spectrum analysis, have not been established, likely due to the highly dynamic HR patterns observed in bats. Further investigation will be essential to define species-specific HRV parameters and to establish reliable physiological markers of autonomic function and emotional states in bats.

## Statements and Declarations

## Acknowledgments

We are grateful to Motoki Kihara, Mayuko Mogi and Daisuke Namazue for their assistance with the recordings. We also thank Yu Teshima and Andreas Mayer for their helpful comments on the additional analysis.

## Ethics statement

All experiments complied with the Principles of Animal Care (publication no.86-23 [revised 1985] of the National Institutes of Health) and all Japanese laws. All procedures were approved by the Animal Experiment Committee at Doshisha University.

## Data accessibility statement

Data are available at figshare (doi: https://doi.org/10.6084/m9.figshare.29614925.v1). Other raw data supporting the findings of this study are available from the corresponding author on a reasonable request.

## Artificial Intelligence (AI) declaration

We have not used AI-assisted technologies in creating this article.

