## Supplementary material for "Mothers respond to biological pup calls with heart rate changes in Japanese house bats, *Pipistrellus abramus*": Table S1

**Table S1. Pup vocalization parameters at 2 and 30 days after birth (DAB).**
Initial frequency (IF), terminal frequency (TF), and duration are shown as mean ± standard deviation values.

| Pup | Age | | | | | |
|---|---|---|---|---|---|---|
| | 2 DAB | | | 30 DAB | | |
| | IF [kHz] | TF [kHz] | Duration [ms] | IF [kHz] | TF [kHz] | Duration [ms] |
| Aa | 30.2±2.0 | 21.6±1.4 | 23.4±5.0 | 69.6±5.6 | 44.2±1.8 | 1.3±0.1 |
| Ab | 35.1±2.1 | 20.7±1.2 | 19.6±6.4 | 76.5±5.8 | 43.3±2.0 | 1.8±0.2 |
| Ac | 33.8±1.0 | 21.3±0.5 | 26.0±2.7 | 58.6±7.5 | 39.5±2.7 | 1.7±0.3 |
| Bb | 35.9±2.0 | 21.8±0.8 | 33.3±5.1 | 80.9±3.6 | 44.0±1.5 | 2.1±0.1 |
| Bc | 35.7±2.2 | 20.6±1.1 | 21.8±4.3 | 86.5±5.0 | 47.9±1.8 | 1.7±0.2 |
| Ca | 36.2±2.1 | 20.4±1.3 | 13.0±3.0 | 70.8±4.1 | 45.9±2.1 | 1.8±0.2 |
| Cb | 32.7±1.2 | 18.3±1.4 | 23.4±4.8 | 58.0±2.5 | 41.1±2.0 | 2.4±0.5 |
| Cc | 33.6±1.7 | 22.5±0.6 | 22.9±3.3 | 71.8±4.6 | 45.8±2.0 | 1.6±0.3 |
| Da | 32.8±2.0 | 20.9±0.8 | 23.6±4.1 | 76.1±2.7 | 42.5±1.4 | 2.0±0.2 |
| Dc | 39.0±2.1 | 23.5±0.6 | 23.1±3.7 | 78.5±2.2 | 46.4±1.3 | 1.5±0.1 |
